# An injectable meta-biomaterial

**DOI:** 10.1101/2020.01.30.926931

**Authors:** A. Béduer, F. Bonini, C. Verheyen, P. Burch, T. Braschler

## Abstract

We present a novel type of injectable biomaterial with an elastic softening transition. The material enables *in-vivo* shaping, followed by induction of 3D stable vascularized tissue adopting the desired shape. We establish the necessary geometrical and physical parameters by extensive numerical simulation. Irregular particle shape dramatically enhances yield strain for *in-vivo* stability against deformation, while friction and porosity provide the elastic softening transition as an emergent meta-material property. Accordingly, we synthesize our injectable meta-biomaterial as a suspension of irregularly fragmented, highly porous sponge-like microgels. The meta-biomaterial exhibits both high yield strain, and the desired novel elastic softening transition for *in-situ* shaping and unprecedented dynamic matching of adipose tissue mechanics. *In vivo*, predetermined shapes can be sculpted manually after subcutaneous injection in mice. The 3D shape is maintained during excellent host tissue integration into the particle pore space. The meta-biomaterial sustains vascularized connective tissue to the end of one-year follow-up.

---

Aging, disease, trauma, congenital deformities and surgical sequelae can all lead to a loss or lack of soft tissue volume, producing a major and increasing medical demand for tissue reconstruction strategies^1–4^. Ideally, such procedures should be minimally invasive to reduce the patient burden and decrease potential surgical complications^5,6^. For soft tissue reconstruction, this requires a tissue bulking agent that can be injected through a thin needle into a target site in the body^4^. However, after deployment this bulking material must maintain a stable three-dimensional (3D) configuration to provide adequate volume for the missing soft tissue^7^. Beyond injectability and volume maintenance, surgeons impose the additional requirement of *in situ* shapeability in order to smoothly match the arbitrary geometry of a patient’s tissue defect^8^. Even if the three conditions of injectability, shapeability and stability are satisfied, this theoretical tri-state material should also exhibit a high degree of biocompatibility while closely matching the local tissue mechanics^9,10^.

In practice, such an ideal material is exceedingly difficult to engineer. For example, fluid-like behavior needed for minimally-invasive delivery through a needle limits mechanical stiffness^11^. As a result, many of the currently available injectable agents are too soft^11^ to match the local tissue properties^12,13^. Though useful for smoothing and enhancing existing shapes, they require improvements to adequately lift and shape three-dimensional tissue volumes^11^. On the other hand, solid-like behavior in preformed implants interferes with *in-situ* shaping, even if minimally invasive delivery is possible by compression or folding^14,15^. This unmet need has launched numerous efforts to engineer injectable, shape-fixable tissue implants, based on various physical or chemical *in-situ* crosslinking mechanisms^16,17^. Emerging strategies include the use of peptide, DNA, or polyelectrolyte and other complexes to impart intrinsic self-healing ability^18–20^. Though such approaches provide improved mechanical properties, they also impose a number of additional chemical constraints and raise concerns regarding biocompatibility and tissue integration^16,21,22^.

Here, we reconcile injectability, shapeability, and long-term volume stability by a radically novel approach: an injectable meta-material^23^. Mechanical meta-materials obtain unusual properties by their mere geometry^23,24^. We hypothesize here that physical interlocking of geometrically designed particles can restore meta-material properties even after transient liquefaction for injection^25–27^. In this novel paradigm, self-healing delivers meta-material physics *in-vivo, in-situ*, in a minimally invasive fashion. We engineer a tri-state microgel meta-biomaterial that combines injectability, solid-state softening for shapeability, and tissue-mimicking mechanical stability^12,13^. The material dramatically increases the accessible stiffness range and for the first time demonstrates a reversible elastic softening transition in an injectable. It exhibits excellent biocompatibility and is colonized to induce formation of vascularized tissue. Taken together, we have successfully integrated the design criteria informed by clinical need to engineer a novel injectable and shapeable biomaterial for stable 3D soft tissue reconstruction.

## Results

### Elastic porous injectable (EPI) biomaterial

The concept of our elastic, porous, and injectable (EPI) biomaterial is illustrated in Figure 1. Under the high shear imposed by injection, the material fluidizes and behaves as a liquid (Fig. 1a). Under intermediate shear, typical of *in situ* shaping and massaging^8,28^, the EPI biomaterial behaves as a weak viscoelastic solid amenable to smooth deformation (Fig. 1b). At rest, the self-healing process is complete and the EPI biomaterial behaves as a soft solid with shear moduli in the lower kPa range to match the native tissue properties^12,13^ (Fig. 1c).

**Fig. 1.**
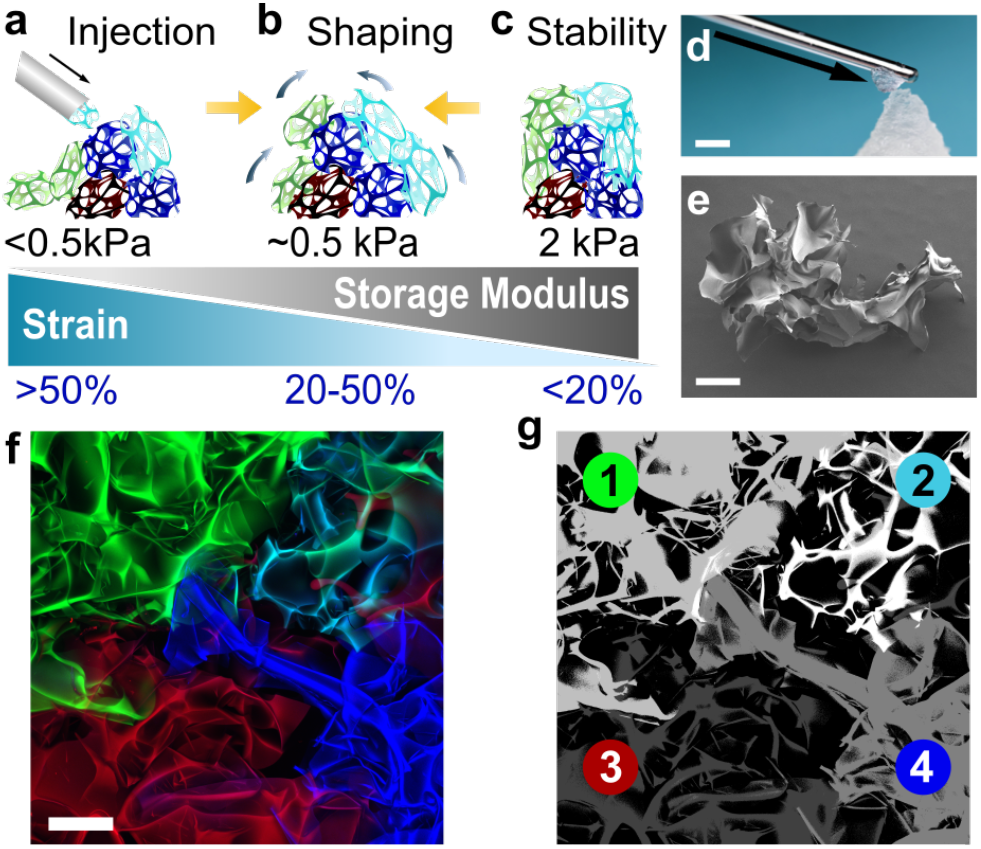
Elastic porous injectable (EPI) biomaterial. **a)** Injectability: Particle liquefaction and mobility under high strain enables minimally-invasive delivery. b) Shapeability: Partial mechanical stability due to geometric particle constraints enables in-situ shaping under intermediate strains. **c)** Volume Stability: Fully interlocked particle state enables long-term three-dimensional stability under low strains. **d)** Macroscopic demonstration of the material properties, depicting fluid-like ejection of particles through a cannula and solid-like 3D shape stability. Scale bar: 4 mm **e)** Scanning electron microscope picture of a single porous particle (brightness proportionally enhanced). **f)** Confocal image showing four interlocking porous particles (maximal intensity z projection). **g)** Particle identity in Fig. 1f, determined by thresholding the individual color channels, and for the doubly labeled particle, co-localization of blue and green after correction for chromatic aberration. Scale bar: 200 μm.

The EPI biomaterial consists of an interlocking suspension of highly irregular, sponge-like microparticles. Its unique material design enables both fluidic injection through a cannula and shapeable three-dimensional stability. A macroscopic demonstration of the behavior is provided in Fig. 1d, while Fig. 1e shows the irregular, porous particle morphology. Fig. 1f and 1g demonstrate microsopic particle interlocking.

### *In-silico* design

To design the EPI biomaterial, outlined in Fig. 1, we first performed a discrete element simulation. This essential step translates the clinical use criteria into a set of design rules that guided our material fabrication strategy. To reiterate, an ideal tissue-filling biomaterial should display injectability, shapeability, 3D volume stability, biocompatibility, and tissue-matching mechanical properties.

A full description of our simulation is provided in Supplementary 1 (mathematical model), 2 (software usage instructions), 3 (implementation details), 4 (test cases), 15 (source code), 16 (API documentation) and 17 (install guide). Briefly, the simulation models the physical interaction between spherical particles by central and frictional forces^25^. To meet the long-standing challenge of symmetrical stress tensor evaluation in granular media^29,30^, we systematically reviewed and corrected the published mathematical framework^25^ and stress tensor evaluation^29^ (Supplementary 1). We then added permanent crosslinks to study both irregular and porous microparticles (Fig. 2a).

**Fig. 2.**
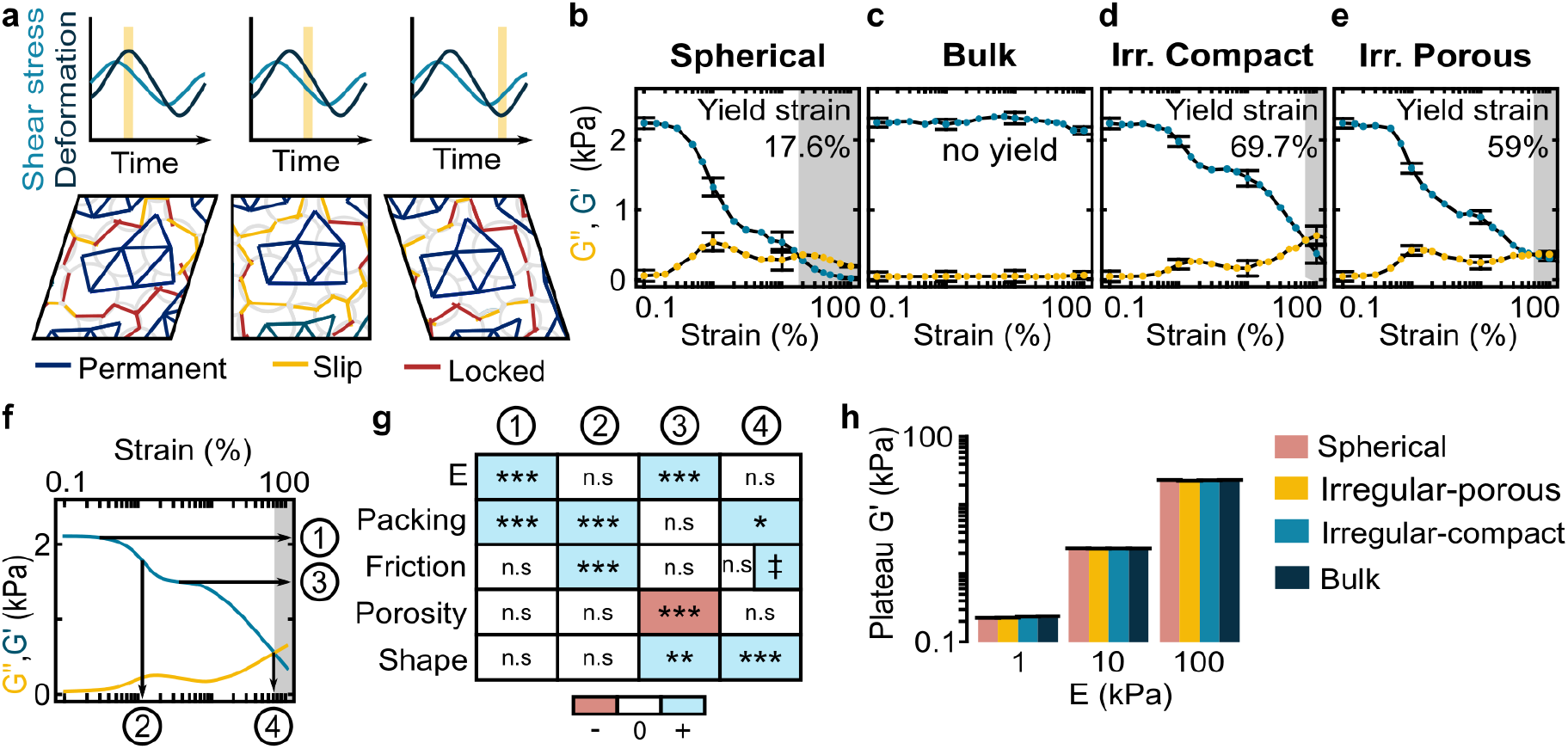
In Silico simulation and design of the EPI biomaterial. **a)** Graphical overview of the simulation. **b)** Elastic storage and viscous loss modulus (G’ and G’’) for a dense suspension of frictional spheres **c)** for a bulk material formed by full cross-linking of every neighboring sphere **d)** for a dense suspension of discrete, compact, irregular particles formed from neighboring spheres **e)** for a dense suspension of discrete, irregular particles with a low crosslink density and free contact interfaces. **f)** Characteristic rheological response with accompanying storage modulus and strain values. ① = low-deformation limit, ② = softening transition, ③ = soft plateau stress, ③ = yield strain. **g)** Overview of the influence of the model parameters on the characteristic values defined in Fig. 2F. ‡ The friction coefficient has a magnitude-dependent effect, see Supplementary 5, Fig. S5-3. **h)** Influence of the Young modulus of the constituent material on the low-strain limit storage modulus for the different particle geometries. Error bars = one standard deviation. n.s. = not significant. Sample size and statistical testing information in Supplementary 14, items 1 to 26.

Analogous to the empirical characterization of viscoelastic agents for soft tissue reconstruction^8^, we performed *in silico* oscillatory shear rheometry by application of time-varying strain(Fig. 2a). This provides an estimate of the elastic stiffness (elastic storage modulus G’) and the ability to deform (viscous loss moduli G’’)^25,31^.

We first simulated four prototypical scenarios of microgel suspensions: a simple suspension of frictional spherical microgels (Fig. 2b, Supplementary video 18), completely crosslinked elastic bulk material (Fig. 2c, Supplementary video 19), compactly crosslinked irregular particles (Fig. 2d, Supplementary video 20) and loosely crosslinked particles with a decreased internal crosslinking density (Fig. 2e; Supplementary video 21). An elastic softening transition was found for the frictional spherical microgel suspension as expected^25^ (Fig. 2b, absent in non-frictional control provided in Supplementary 5). But the yield strain was substantially below the 50% typically required for a soft tissue filler^8^ to withstand physiological movement^32^. On the other hand, bulk crosslinking (Fig. 2c) abolished the yielding transition altogether. Irregular particles finally (Fig. 2d) exhibited yield strain well above 50%, enabling both injectability and *in vivo* shape stability. This establishes our first design rule: The material should be a suspension of irregular rather than spherical particles.

Quantitatively, the elastic softening transition was better preserved in loosely (vs. densely) crosslinked particles (Fig. 2e). The softening behavior is critical both for shapeability and for matching the mechanical properties of local adipose tissue, which also demonstrate strain softening under shear^12,13^. We thus obtain our second design rule: the particles should have a low density of internal crosslinks with frictional intra-particle porosity.

With our first two design rules established, we generalized the desired rheological behavior (Fig. 2f) and performed systematic analysis of the model’s parameters (Fig. 2g, Supplementary 5). The analysis confirmed that the geometric parameters (particle shape and porosity) dominate the high-strain behaviors^33^ of yielding (④ in Fig. 2f and 2g) and the soft plateau (③), important for shaping and injection. Conversely, the mechanical parameters (friction, packing, and Young’s modulus) dominate the low-strain behaviors of softening (②) and the low-strain plateau shear modulus (①), important for tissue interaction.

Specifically, Fig. 2h indicates that the low-strain storage modulus was proportional to the Young’s modulus of the constituent microgel material (linear regression, P=8*10^−88^), but independent of the particle geometry (P=0.50) or crosslinking density (P=0.29) (statistical analysis in Supplementary 14, item 27). Ideally, our material’s low-strain mechanical behavior would match that of native soft tissue (lower kPa range^12,13^). Thus we obtain our third design rule: the bulk precursor from which we synthesize our microparticle suspension should have a storage modulus close to the low strain limit of adipose tissue.

The first three design criteria produce the optimal rheology for injectability, shapeability, and tissue-matching 3D stability. Our last guideline comes from the established rule that scaffolds should have pores of at least 50 μm diameter to ensure vascularization and colonization^34,35^. Thus we obtain our fourth and final design rule: the microparticles should have a mean pore size of at least 50 microns.

### Synthesis and mechanical characterization

Our *in silico* analysis revealed that an injectable, shapeable, and volume-stable material could be achieved by a densely packed suspension of elastic, frictional microparticles with irregular porous geometry (Fig. 2). To further ensure biocompatibility we based our synthesis on the cryogelation^36^ of carboxymethylcellulose^34,37^. The polymer content was adjusted to obtain a porous bulk material with an elastic storage modulus (G’) of 2.4kPa +/− 0.9kPa (Supplementary 6). We therefore satisfied design rule #3, which stated that the bulk precursor elastic storage modulus should be in the low-strain limit of adipose tissue^12,13^.

Through forceful extrusion we then fragmented the highly porous bulk material^37^ into irregular microparticles with a diameter of 805 +/− 363 μm (Supplementary 7). Thanks to the intra-particle porosity, the pore space in the resulting EPI biomaterial displays an intricate and widely connected morphology (Fig. 3a). This unique porosity contrasts starkly with that of a traditional microgel suspension like the chromatography medium Sephacryl S200^38^, where pores are limited to the interstices between the dense spherical particles (Fig. 3b). Therefore the abundance of large frictional pore spaces fulfilled design rules #2 and #4, while the exceptional irregularity fulfilled design rule #1. Thus we obtained an elastic, porous, and injectable (EPI) biomaterial that satisfied our pre-defined design criteria.

**Fig. 3.**
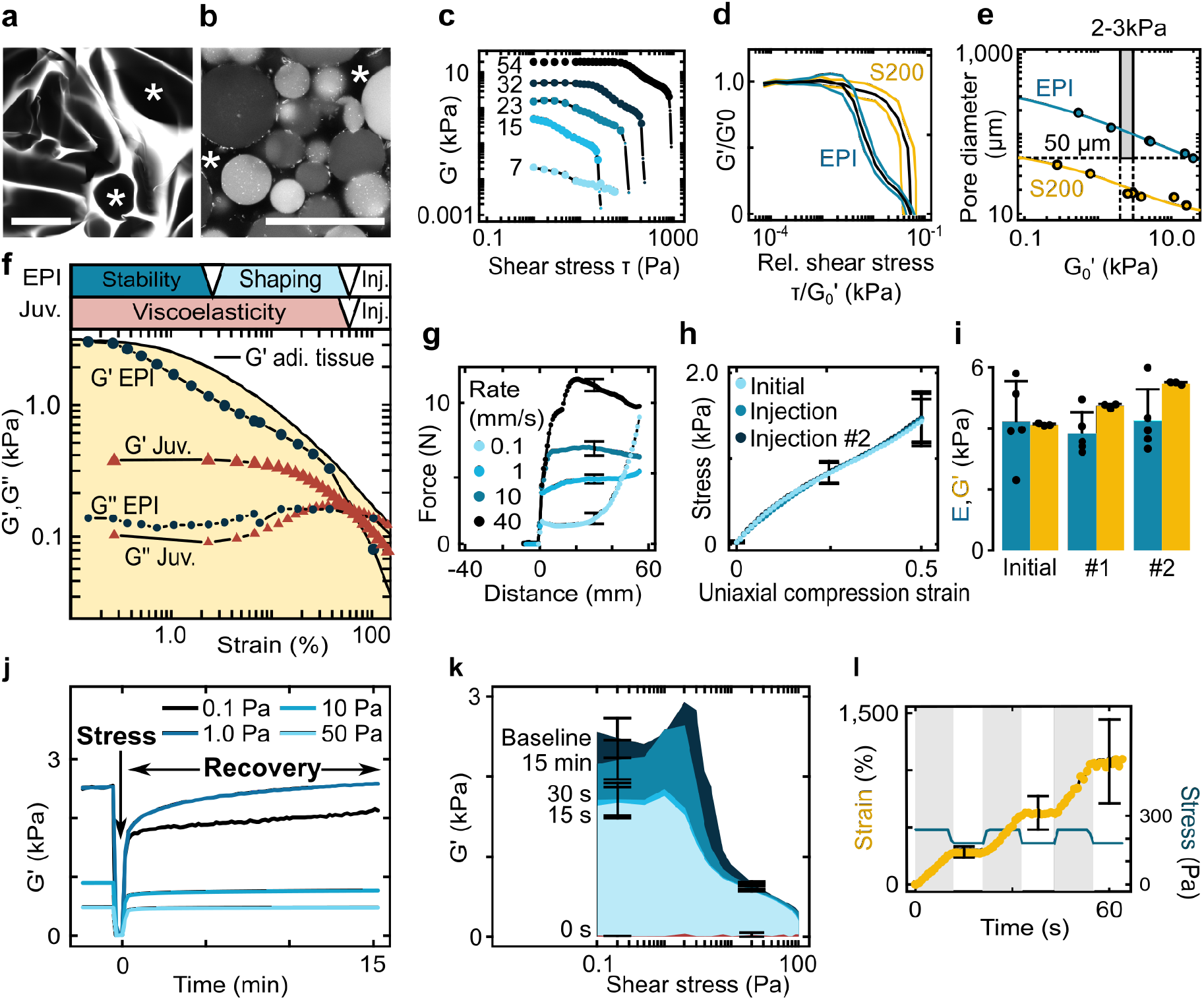
In vitro characterization of the EPI Biomaterial. Confocal images of a) the EPI biomaterial (stained with rhodamine 6G) and b) Sephacryl S200 (autofluorescence). The stars denote pore space. Scale bars = 200 microns. **c)** G’ of the EPI biomaterial as a function of applied oscillatory stress for various EPI biomaterial polymer concentrations (indicated, mg/ml). Large symbols (⚫): solid-like behavior (G’>G’’); small symbols (•): liquid like behavior (G’’<G’). Each curve represents a single measurement. **d)** Normalized master curves for EPI and Sephacryl 200, error lines +/− one standard deviation. **e)** Pore size vs. low-stress G_0_’ values. Grey box (▭): 50μm minimal pore size and a G_0_’ value between 2 and 3kPa required for matching adipose tissue^12,13,34,35^. **f)** Comparison of the EPI biomaterial (26mg/mL) to commercial dermal filler Juvederm Voluma (single samples) and re-plotted literature G’ data on bovine retro-orbital adipose tissue^13^ **g)** EPI biomaterial injectability. **h)** Uniaxial compression analysis of a sample before passage through the cannula, after the first passage, and after two consecutive passages. **i)** Comparison of E and G_0_’ before and after 1 or 2 passages through the cannula. **j)** Recovery of the G’ values after a period of liquefying shear (“Stress”). **k)** Level of G’ recovered after different time-points as a function of continuous oscillatory shear stress (same data set as for Fig. 3j). **l)** Rapid and reversible self-healing by transition between liquid state and solid state. Error bars = one standard deviation. Sample sizes and statistical testing details in Supplementary 14, items 28-41.

With the fabrication phase complete we began extensive physical characterization. We first investigated the presence of an elastic softening transition, along with high yield strain, as predicted by our simulation (Fig. 2). For this, we subjected the EPI biomaterial (at various levels of polymer concentration) to increasing oscillatory shear stress (Fig. 3c, Supplementary 10). We indeed observed a stable elastic plateau, a unique softening transition, and yielding at high strain (62 +/− 5%) for a wide range of polymer concentrations, Supplementary 10). Normalization to the low-strain plateau value (G_0_’)^39^ confirms that these features are conserved for the EPI biomaterial across all tested concentrations (Fig. 3d). As expected for an essentially non-frictional^33^ micro-hydrogel suspension (Supplementary 5, ref. ^25^), no distinct softening behavior was seen with the Sephacryl S200 material (P=1.7*10^−8^ for difference at 1% strain, item 29 Supplementary 14). Further, in agreement with our numerical results for spherical particles, the yield strain was significantly lower (24 +/− 6%, P=4.8*10^−6^ vs. EPI biomaterial, Supplementary 10 and item 27 in Supplementary 14). We therefore succeeded in engineering a novel biomaterial that displays not only elastic behavior with high yielding, but also a new softening transition that should enable both shaping and tissue matching.

Our next aim was precise mechanical matching to adipose tissue^12,13^, while conserving a pore size greater than 50μm required for vascularization^34,35^. Fig. 3e demonstrates indeed an inverse relation between G_0_’ and the average pore diameter: Fluid withdrawal to increase polymer concentrations lowered both total pore fraction and average pore size (Supplementary 8). With the EPI biomaterial, adipose tissue stiffness G_0_’=2−3kPa^12,13^ was matched while successfully maintaining a pore size of 100-120 μm (Fig. 3e, Supplementary 8 and 9). On the contrary, sufficient stiffness in the Sephacryl S200 material could only be achieved at insufficient pore size, and vice versa. The EPI biomaterial design thus specifically enables the joint fulfillment of both mechanical and geometric requirements.

We further assessed dynamic adipose tissue matching by rheological comparison of the EPI biomaterial to the published rheological response of adipose tissue (bovine, retroorbital^13^). We find that the EPI biomaterial dynamically mimicks the overall adipose tissue response over a wide range of deformations (Fig. 3f). In addition, the EPI biomaterial shows yielding behavior similar to the established dermal filler Juvéderm Voluma (Fig. 3f). The Juvéderm Voluma filler is known to consist of irregularly shaped microparticles^40^, confirming the predicted importance of this feature for high yield strain (57+/−7 % for Juvéderm Voluma®, P=1.0 vs. EPI biomaterial, Supplementary 10). The softening transition engineered into the EPI biomaterial additionally offers the unprecedented dynamic tissue matching.

For minimally invasive delivery, the EPI biomaterial must be injectable. Fig. 3g (controls in Supplementary 11) shows that the forces required for extrusion of the EPI are well below the upper bound of approximately 20N reported for facile manual injectability in a clinical setting^20^. Fig. 3h further indicates conservation of material properties throughout the injection process, as the uniaxial compression response was identical after 1 or 2 injection cycles compared to the original compression response. The Young modulus (E’) was conserved over subsequent injection cycles (Fig. 3i, P=1.0, item 35 in Supplementary 14), whereas the elastic storage modulus G’ actually slightly increased (Fig. 3i, P=3.4*10^−8^, item 36 in Supplementary 14). This increase was likely due to dehydration during transfer steps. Overall, we conclude that the EPI biomaterial can easily be injected and retains its mechanical properties after minimally invasive delivery through a cannula.

Finally, we investigated the physical self-healing process enabling restoration of solid-like mechanical integrity after the injection process. For Fig. 3j, self-healing of the EPI biomaterial following fluidization was assessed under weak (0.1Pa) to strong (50 Pa) continuous oscillatory stress. Self-healing was efficient for all stress levels, but different time-scales were involved. For smaller shear stresses, substantially higher G’ values were eventually achieved, but the final value was reached on a time scale of 10 minutes or above (Fig. 3j, Fig. 3k, Supplementary 12). For strong continuous shear stress, relatively low but stable G’ values were rapidly established. Using an abrupt switch of stress slightly above and below yield stress, we could indeed demonstrate near-instantaneous recovery (Fig. 3i, additional creep-recovery data in Supplementary 13). We believe that the immediate self-healing after injection could stabilize the injected implant to prevent unwanted spreading, followed by attainment of a much higher elastic storage modulus to better match the local adipose tissue at longer time scales.

In summary, we have engineered an elastic, porous, and injectable (EPI) meta-material (Fig. 1) according to the design specifications derived from clinical criteria and *in silico* simulation (Fig. 2). The EPI biomaterial displays a unique tri-phasic rheology: Yield stress enables facile manual injection, followed by rapid initial self-healing to prevent spreading. The reversible elastic softening transition then progressively recovers an elastic storage modulus of 2-3kPa, in line with that of the local soft tissue^12,13^. That same softening transition enables tissue-matching across a wide range of applied forces.

### In-vivo

After the design, synthesis, and extensive *in vitro* testing of our engineered meta-material, we then investigated *in vivo* performance of the EPI biomaterial. With the ultimate goal of clinical implementation, we specifically examined minimally-invasive delivery, *in situ* shapeability, long-term 3D shape maintenance and tissue integration.

For this, we manually injected EPI biomaterial into the subcutaneous space of CD1 mice and then shaped it by application of gentle external force (Fig. 4a). We found that fluidization enabled facile delivery through a 20G catheter. Magnetic Resonance Imaging (MRI) confirmed that the material behaved as a well-defined, cohesive implant in the dermal space (Fig. 4b).

**Fig. 4.**
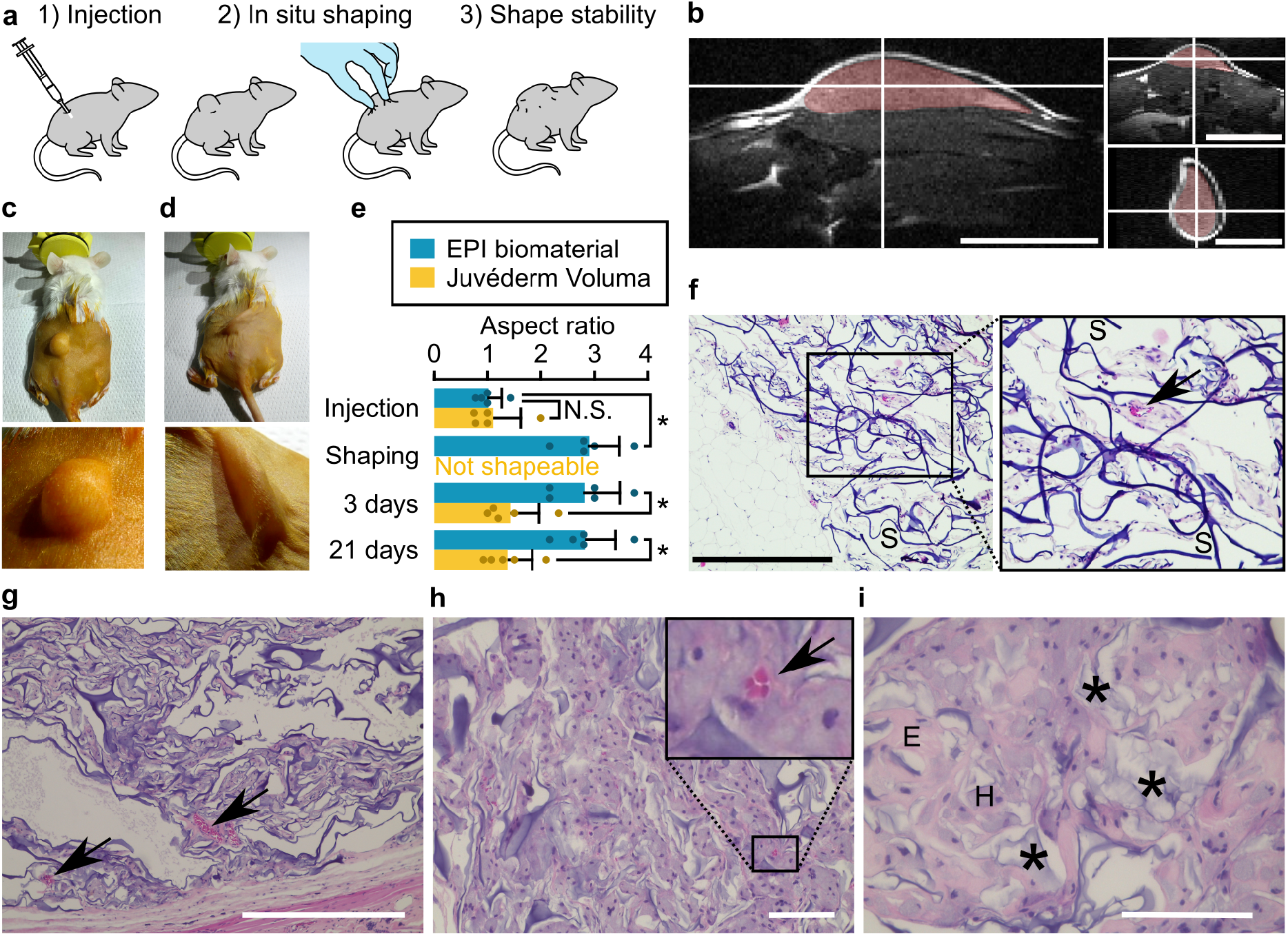
In Vivo Application of the EPI Biomaterial. **a)** Experimental workflow **b)** MRI imaging of the EPI implantation, EPI labeled in false color (center of injected material at hairline crossing). Scale bars = 1mm. **c)** A 400μL bolus of the EPI biomaterial after minimally-invasive injection. **d)** EPI arbitrary 3D in situ shaping by application of moderate external forces. **e)** Quantification of the maintenance of the shape by the aspect ratio of length along the injection direction to width of the implant for EPI and Juvéderm Voluma®. **f)** Histology of the EPI after 21 days of implantation, EPI scaffold material in dark purple (S). Towards the left lower corner: endogeneous adipose tissue. Scale bar=500 μm. **g)** Histology of the EPI at 6 months. Scale bar=500 μm. Not shaped, 200μL bolus. **h)** Histology of the EPI at decreased synthesis concentration to accelerate degradation, at 1 year. Example of a small vessel shown in magnified inset. Scale bar=100μm. Not shaped, 200μL bolus. **i)** Same condition as 4h, higher magnification image. Degrading scaffold denoted by stars, E=example of eosinophilic region, H=example of region stained by hematoxylin. Images 4h and 4i were adjusted regarding white balance and luminosity on cell-free areas for direct comparison to 4g. Error bars = one standard deviation. Vasculature outlined by arrows. Statistical testing details and sample in Supplementary 14, items 43-46.

During a time window of about 20 minutes following bolus injection, the material could be re-shaped *in situ* to produce a new desired shape, retained spontaneously (Fig. 4c, 4d, 4e, Video Supplementary 22). This indicates our success in engineering a shapeable material that provides substantial lifting capacity for 3D soft tissue reconstruction. For comparison we also injected a bolus of the hyaluronic acid dermal filler (Juvéderm Voluma®) but were unable to perform *in situ* shaping, since the material would spread rather than adopt a new shape.

After the shaping time window, the shape stabilized and excessive force was required to dislodge the mechanically interlocked particles. We assessed the long-term stability by attempting to reshape the 3D volume (by massaging) every day during the first week, and then once a week for the remainder of the 3 week follow-up. Despite this gentle mechanical challenge, the shape remained remarkably stable (Fig. 4e).

On the time scale of hours to days, initial stabilization by mechanical self-healing is most likely relayed by irreversible biological processes such as fibrin coagulation^41^, and progressive tissue ingrowth (Fig. 4f, 21 days). At 6 months (Fig. 4g, not shaped), we indeed find most of the implant pore space colonized, with an onset of scaffold degradation. We then assessed bio-integration at the scale of 1 year in Fig. 4h and 4i (reaction mix diluted 2:1 with deionized water to enhance our chances to observe late biodegradation stages, not shaped). We find areas of advanced EPI biomaterial degradation at one year (stars in Fig. 4i). A fibrovascular tissue (pink in Fig. 4h and 4i, example labeled “E” in Fig. 4i) nevertheless persists, along with presumably phagocytic cells processing the degrading material (light purple in Fig. 4h and Fig. 4i, example labeled “H”). Detailed quantification of long-term volume stability, colonization and biocompatibility of the EPI biomaterial in comparison with commercial injectables is to be published elsewhere (A.B., M. Genta, N. Kunz, P. B., T. B., in preparation).

Taken together, our *in vivo* experimentation demonstrated that the unique rheological and morphological properties of the EPI biomaterial translated into a novel capacity to inject, shape, and stabilize customized 3D tissue volumes for tissue induction. Given the range of conditions driving a pathological loss of soft tissue volume, we propose that our new meta-material approach could have a major impact on clinical 3D tissue reconstruction.

## Discussion

We present the design, synthesis, and testing of an *in-vivo* 3D shape tissue engineering material baed on meta-material physics.^23,24^ Numerical simulation allowed us to translate the unmet clinical need of a shapeable soft tissue implant^8,11,16^ into a set of engineer-able mechanical and geometric parameters. With the elastic, porous, and injectable (EPI) biomaterial, we achieved desired shape-stable, softening, and yielding behaviors, along with substantial porosity and a tissue-matching mechanical response. The result was excellent *in-vivo* injectability, shapeability, shape stabilization as well as long-term 3D tissue integration.

In this work we engineered and exploited the elastic softening transition to combine injectability with the full matching of adipose tissue rheology over a wide range of strains. Such an approach allows for a minimally-invasive filler with unprecedented *in vivo* 3D lifting^11^ and shaping capacity. Strain softening is common in ductile materials (like plastics or metal alloys) but it is generally accompanied by large-scale plastic deformation^42^. On the contrary, reversible strain softening with negligible deformation is rare, but has been described previously in actin or cellulose hydrogels^26,27^. Thought to be caused by a buckling of constituent fibers, this response allows the network to maintain rigidity at near static conditions but smoothly and reversibly respond to moderate forces by controlled deformation after softening^26^. Reversible strain softening in microgel suspensions has been conjectured^25,43^, here we provide experimental proof.

Taught by the numerical simulation, the key to this achievement is the combination of porosity and irregular particle shape. Indeed, spherical cryogel particles are reported to display a low yield strain and complete absence of a softening regime^44^, while dermal fillers with irregular hyaluronic acid particles^40^ have high yield strain^8^, but do not display reversible softening.

Our focus on geometrical design of the microstructure characterizes the EPI biomaterial as a mechanical meta-material^24^. Here we employed carboxymethylcellulose-based scaffold chemistry for its known biocompatibility^34^ and obtained excellent bio-integration up to 1 year. Nevertheless, our meta-material design suggests that other scaffold chemistries could be used to realize similar materials. Thus injectable, shapeable, porous materials could be applied to fields as diverse as bone engineering, wound repair, and regenerative organ engineering. By modulating the stiffness, degradation, and biological activity or cell delivery^14,37^, we believe our approach is well suited for a wide range of customized tissue engineering applications beyond 3D soft tissue reconstruction.

## Acknowledgments

Particular thanks go to Aleksandra Filippova for help with team coordination and task scheduling and Daniel Lyubenov for initial help with the simulation. We would further like to thank Benoit Desboilles for SEM imaging, Nicolas Kunz for MRI imaging, and Sidi Bencherif for teaching and initial help with the cryogel technology, Martina Genta and Mariana Martins for help with animal experiments, and Prof. Paul Bowen for letting us use his rheometer. We also acknowledge the support by the following facilities and their staff: microscopic imaging facilities of the University of Geneva (Bioimaging Core Facility of the Faculty of Medicine) and EPFL Lausanne (Bioimaging and Optics Platform), the histology core facility at EPFL and particularly Jessica Sordet-Dessimoz, the Center of Micronanotechnology CMI and the CIBM facility at EPFL, including its animal facility. The simulations were performed on the Baobab Cluster of the University of Geneva. Funding for this work was provided by the Swiss National Science Foundation (PP00P2_163684 and PZ00P2_161347), the Gebert-Rüf foundation (GRS-0043/15), and a Whitaker International Fellowship.

## Author contributions

A. Béduer, F. Bonini, C. Verheyen and T. Braschler contributed to the design of this study. All authors contributed to the writing and proofing of the manuscript. A. Béduer carried out and analyzed the *in-vivo* study. F. Bonini provided the morphological biomaterial characterization and the design of the figures in the manuscript. C. Verheyen performed the mechanical characterization, and T. Braschler wrote the software, ran and analyzed the simulations. P. Burch developed and optimized biomaterial fabrication.

## Data availability

Figures 2, 3 and 4 have associated quantitative raw data. The raw data and the scripts to produce the quantitative figures are provided at the Zenodo repository (https://zenodo.org/record/2653804#.XNBXFIS7Ldk). Qualitative raw data (histology, MRI) is available from the authors upon request.

## Code availability

The full simulation source code is available for download (Supplementary 15). Details of the mathematical model (Supplementary 1), full usage instructions (Supplementary 2), implementation details including pseudocode (Supplementary 3), an automatically generated full API documentation (Supplementary 16) and unit test cases (Supplementary 4) are also provided. A quick installation guide is available in Supplementary 17. The source code is also included into the Zenodo repository (https://zenodo.org/record/2653804#.XNBXFIS7Ldk) for appropriate versioning.

## Online Methods

### Simulation

The simulation was implemented as a custom Python library. It implements the physical interaction framework defined by Otsuki at al.^25^, corresponding to central viscoelastic interaction and tangential friction between spherical particles. To ensure exact rather than approximate conservation of angular momentum, we adjusted the expression of the torque resulting from frictional interaction (Supplementary 1, eq. S1-3).

To correctly evaluate stress tensors at large amplitudes, we implemented the stress tensor evaluation framework provided by Nicot et al.^29^. As unit testing (Supplementary 4) revealed non-symmetric stress tensors in some cases, we re-evaluated the calculations by Nicot et al. ^29^. We found a probable integration mistake regarding the inertia matrix for the contribution of unbalanced torques to the stress tensor (Supplementary 1). This directly concerns eq. 29 in ref. ^29^, and as result also eq. 30-35 in ref. ^29^; we use a corrected version of eq. 34 in Nicot et al.^29^, supplied as eq. S1-24 in Supplementary 1. To avoid complete interpenetration of neighboring particles at large shear, we added a non-linear term to the repulsive elastic interaction. We ensured that in the small compression limit, we recover the linear repulsion law used by Otsuki et al.^25^ (Supplementary 1, eq. S5b).

Finally, we also implement the possibility of permanent crosslinks between neighboring spheres. The permanent crosslinks remain intact regardless of the geometrical separation of the spheres, generating attractive forces if the spheres are separated beyond touching distance. They also do not allow for frictional slippage. Additional mathematical details of the implementation are given in Supplementary 1. The source code can be downloaded as Supplementary 15; software usage, implementation details and test cases are provided respectively in Supplementary 2,3 and 4. The full API documentation is available as Supplementary 16, and quick install instructions as Supplementary 17, simulation videos are provided as Supplementary 18 – Supplementary 21.

The simulations were run on the Baobab cluster of the University of Geneva. Instructions on how to replicate our simulations and data analysis are provided in Supplementary 2.

### Statistics

Statistical evaluation and graphing were done using the R-Cran free software, version 3.2.3^45^. Central tendency was reported by arithmetic mean values, and variability by indication of single standard deviations. For comparisons with 5 or less measurements per group, individual values were also graphed (Fig. 3i, Fig. 4e). Linear regression was used to assess continuous effects, and paired or unpaired, Student (t) tests were used to compare individual conditions, with P-values reported after Bonferroni multiple testing correction^46^ for all tests performed per subfigure. For all tests, the Shapiro-Wilks test was used to assess normality of the residuals. If significant deviation was found, an alternative test was evaluated: generalized linear models with identity link but expection-value dependent variance (quasi-likelihood estimators^47^) to accommodate heteroscedasticity in linear regression, and Wilcoxon signed rank (paired) or Mann-Whitney U (unpaired) for binary comparison. All tests performed are reported in Supplementary 14, including their effect sizes and normality testing results.

For the statistical evaluation of the characteristic strains and stresses of the simulated elastic storage modulus and viscous loss modulus curves (G’ and G”) as a function of the model parameters (Fig. 2G) a bootstrapping approach was taken^48^. For a given set of model parameters, 5 subsets among the simulations carried out for the various strain amplitudes are drawn randomly with a 20% probability for a given simulation to be included. For each subset, slightly different estimations for characteristic stresses and strains as defined in Fig. 2f are thus obtained. By linear regression of these values against the physical parameter being varied, the P-values depicted in Fig. 2g are obtained after Bonferroni correction^46^. To reduce dependence of the P-value on the random draw, we averaged the underlying F-statistics over 500 bootstrapping runs^48^.

### Reagents

1,4-Piperazinediethanesulfonic acid (PIPES), adipic acid anhydride (ADH), 1-ethyl-3-(-(3-dimethylaminopropyl)carbodiimide (EDC), rhodamine 6G hydrochloride, 4′,6-diamidino-2-phenylindole (DAPI), 6-aminofluorescein, 37 % HCl solution, NaOH pellets and Sephacryl S200 HR were all purchased from Sigma-Aldrich. Carboxymethyl cellulose (AQUALON CMC 7LF PH, 90.5 KDa, DS: 0.84) was purchased from Ashland. EDTA Solution (0.5 M, pH 8.0) was obtained from ThermoFisher Scientific. Saline solution (0.9% NaCl) was purchased from Bichsel. Disposable sterile vacuum filtration systems (pore size 0.2 um, filter capacity 1000 mL) were purchased from Sigma-Aldrich, whereas cell strainers (40 μm) were obtained from VWR.

### Biomaterial synthesis

Cryogel scaffolds were synthesized as reported^37^, with minor modifications. Briefly, carboxymethyl cellulose (13.56 g), PIPES Buffer (6.30 g), adipic acid anhydride (486 mg) and NaOH pellets (1.20 g) was dissolved to a final volume of 1000 mL in deionized water and the solution filtered at 0.2 μm. This solution could be stored in a 4°C for up to a month prior to use in a fridge. Then EDC (2.70 g) was added. After mixing, the solution was filled into 35 mL plastic syringes. The syringes were closed with a syringe cap and then placed into a freezer set to −20°C.

After 2 days the syringes were removed from the freezer and allowed to reach room temperature. The EPI biomaterial was obtained by forceful extrusion through a 22G catheter. On a filter system, the biomaterial was washed with 10mM EDTA, followed by incubation with 2 M NaOH solution (2h), washing with physiological saline (3x). For *in-vivo* experiments the biomaterial was sterilized for 20 min at 121°C.

### Microscopic imaging

Confocal images were captured on a confocal Zeiss LSM 700 driven by Zen 2010b version service pack 1 (Zeiss), using a 10x Plan Neofluar lens with a numerical aperture (NA) of 0.3 or a 20x PlanApochromat objective with NA=0.8. Alternatively, we also used a Zeiss LSM 800 driven by ZEN 2.3 with ZEN module Tiles (Zeiss), also using either 10x PlanApochromat lens with NA=0.45 or a 20x PlanApochromat with NA=0.8. Images of histological slides were acquired with a Leica DM750, with achromat lenses of 10x (HI PLAN, NA=0.25), 20x (HI PLAN, NA=0.4) or 40x magnification (HI PLAN, NA=0.65), with image storage directly to an SD card by a Leica ICC50 HD Camera without additional software. Table 1 below details the microscope setup used for each experiment or figure.

**Table 1.** Microscopic imaging. Summary list of the microscopes and objectives used. Further details on the microscopes and lenses in the text.

| Figure | Microscope / Objective | Adjustments |
| --- | --- | --- |
| 1f | Zeiss LSM 700, 10x | No adjustment |
| 3a | Zeiss LSM 700, 10x | No adjustment |
| 3b | Zeiss LSM 800, 20x | No adjustment |
| 4f | Leica DM750, 10x | No adjustment |
| 4g | Leica DM750, 10x | No adjustment |
| 4h | Leica DM750, 20x | Adjustment of white balance and luminosity on cell-free areas to match Fig. 4g |
| 4i | Leica DM750, 40x | Adjustment of white balance and luminosity on cell-free areas to match Fig. 4g |

For visualization and morphological quantification, the EPI biomaterial was stained with 5microgram/mL Rhodamine 6G hydrochloride (Fig. 1 and 3a) or 4′,6-diamidino-2-phenylindole (DAPI, by affinity to the final material) or 6-aminofluorescein (by inclusion of 10 micromolar aminofluorescein into the synthesis mixture^37^), or DAPI and aminofluorescein (Fig. 1). For the reference material, Sephacryl S200, auto-fluorescence images (excitation 353nm, emission 405nm and above) was used for visualization and quantification.

Scanning electron microscope SEM images were finally acquired with a Zeiss Merlin SEM, equipped with a Gemini II column and the ZeissSmartSEM acquisition software.

### Pore size and pore fraction

After staining (for the EPI biomaterial) and confocal image acquisition, Fiji software^49^ (ImageJ version 2.0.0-rc-44/1.50g) was used to visualize z-stack images, and to quantify pore size and fraction as well as particle size. For pore fraction quantification, images were acquired at a resolution so that several pores fit across the image, implying generally the use of a 10x objective for the EPI biomaterial vs. a 20x objective for the Sephacryl S200 control material. The EPI biomaterial was stained with rhodamine 6G (excited at 550 nm) whereas autofluorescence images (excited at 355nm,) were acquired for the Sephacryl S200 reference. The images were thresholded automatically using the Li algorithm^50^, built-in in Fiji. In case of evident gross misinterpretation of the images by the algorithm, thresholding was carried out manually instead. The wall and pore fraction could then be estimated from the fraction of white and black pixels; for pore size estimation, the maximal sphere fitting algorithm by Beat Münch et al.^51,52^ was used (Supplementary 8). For each polymer concentration and material, between 20 and 34 images were quantified.

Particle size distribution (Supplementary 7) finally was evaluated from large tile-stitched images acquired on dilute particle suspensions using a Zeiss LSM 800 with a 5x PlanApochromat objective, NA=0.16, after staining with rhodamine 6G as for pore size. Standard Fiji^49^ routines were then used on the resulting large images: binarization by manual thresholding, digital filling of holes, and particle size analysis (minimal size 10 micrometers^2^, circularity at least 0.1).

For all confocal imaging, observation chambers avoiding both evaporation and mechanical compression were used. These chambers consisted of a 1.5mm high Perspex sheet (Evonik) cut to the shape of a microscope slide (75mm × 25mm) using a laser cutter (HobbyLaser, FullSpectrum Engineering, commandeered by FullSpectrum Laser RetinaEngrave3D, Version 4.423). A rectangular sample reservoir (30mm × 12.5mm) was further cut into these Perspex microscope slides. A microscope coverslide was glued permanently to the lower side of the Perspex microscope slide using bathroom silicone glue (Coop BricoLoisir, Switzerland). After filling with the sample to be observed by confocal microscopy, the observation chamber was closed by capillarity using a second microscope coverslide.

### Rheology

Rheological measurements were carried out on a Haake Rheostress RS100 5Ncm apparatus, using the RheoWin software (RheoWin Job Manager: version 3.61.0005) to control the apparatus and acquire data. To avoid wall slipping, roughened surfaces were obtained by gluing a rough cleaning cloth (Miobrill, Migros Switzerland, ref. 7065.206 / 15.02.2330) with hot glue (UHU, LT110, local hardware store) to the surfaces in contact with the material. Experiments were generally conducted in a custom cup geometry (Supplementary 10), except for where the sample volume was too low (Juvéderm Voluma sample for Fig. 3f and the injected samples for Fig. 3i, measured with plate-plate geometry: Haake PP20, ref. 222-0586). In addition, we measured repeated solid-liquid transition (Fig. 3l) with a vane geometry (Haake FL-16, ref. 222-1326) to limit sample movement. Further details and comparison of the different geometries are given in Supplementary 10. A solvent trap (Haake ref. 222-0607) was used whenever possible to limit sample evaporation.

With the exception of the self-healing experiments, we generally preconditioned the material with an oscillatory stress sweep from 1-10 Pa and back at 0.2Hz before applying the desired shear protocols to minimize the impact of shear history. In rare cases of very dilute material, we found this to induce significant shear softening or even yielding, in which case we changed to a preconditioning sweep from 0.1 Pa to 1Pa and back, still at 0.2Hz. After data acquisition, we exported the data to text files (RheoWin Data Manger: version 3.61.0005) and completed data treatment and graphing in R-Cran (version 3.2.3).

Rheological master curves represent the elastic modulus G’ normalized to the plateau value at low stress^39^. They are obtained by normalizing G’ curves with respect to the low-strain G_0_’ plateau value^39^. We evaluate the low-strain limit G_0_’ as the average of the G’ values for the measurement points with low applied shear stress (τ<2Pa). We further take care to exclude points showing already an onset of softening or liquefaction by also imposing G’’(τ)<0.1*G’(τ). We also normalize the applied stress τ to G_0_’. After visual verification that indeed all the different G’ curves obtained at different polymer concentrations collapse onto a single master curve after normalization of both the G’ and τ, we obtain the main master curve by averaging of the normalized curves, along with evaluation of the standard deviation. For the more horizontal part of the master curve (G’/G_0_’>0.5 for the Sephacryl S200 master curve, G’/G_0_’>0.1 for the EPI biomaterial) we perform vertical averaging of the G’/G_0_’ associated with a given interval of applied stress τ, whereas for the steepest part of the master curves, we rather average the τ associated with a given interval of G’/G_0_’ values.

### Uniaxial compression and injectability testing

We performed uniaxial compression testing on a TextureAnalyzer TA.XT plus machine by Stable Microsystems, using the TestMaker program (Version 4.0.6.0) supplied by the manufacturer to program the tests, and the TextureExponent 32 program (Version 4.0.13.0), also supplied by the manufacturer, to run them. After acquisition, the data was exported as text and analyzed using R-Cran.

For compression analysis of the elastic porous injectable (EPI) biomaterial, a disk of 20mm diameter and approximately 6mm height was shaped under a 20mm diameter chuck. The chuck was then moved by feedback to the zero force condition. This defined the original sample height, usually in the range between 5 and 6mm depending on the actual amount of sample. From this position, compression by 50% of the height at a speed of 0.01mm/s was then carried out, acquiring the force and position data. The force was low-pass filtered to reduce noice, and was then converted to stress by dividing through the contact area (circle of 10mm radius), whereas deformation was expressed as deformation strain relative to original sample height.

For injectability analysis, a 1mL syringe (BD) was loaded with test material (EPI biomaterial, deionized water, or air), and equipped with a blunt delivery cannula (Thiebaud Biomedical devices, ref. F9020100, outer diameter 2mm, length 10cm). The syringe was then placed on custom holder. The piston was then moved by using the TextureAnalyzer XT Plus machine at the desired rate, while recording force and distance.

### Animal experiments

All *in-vivo* experiments were approved by the Animal Care and Use Committee of the Canton of Vaud, Switzerland (Authorization VD 3063). Female, adult CD1 mice between 12 and 20 weeks of age were obtained from Charles River (Bar Harbor, Maine, USA) and allowed to acclimatize in the animal facility for at least 1 week prior to implantation. Room temperature was kept at 22+/−2°C with 12 hours light/dark cycle and normal diet *ad libitum*.

Prior to injection, animals were anesthetized with 4% (2% for maintenance) isoflurane (Animalcare Ltd), using an ophthalmic gel (Viscotears, Alcon) for eye protection. The area for injection was shaved and disinfected with betadine (Mundipharma Medical Company). For injection, a small access was created in the skin with a 18G needle, followed by injection of biomaterial samples (2 injection sites per animal, or max. 400 μL into single site) through a 20G catheter (BD Biosciences). No sutures were required. Animals were monitored weekly throughout the study.

When shaping was first attempted with the Juvéderm Voluma®, the material was found to flow and redistribute during the procedure, and during follow-up, we observed large swelling responses over the following weeks. To avoid unnecessary strain on the animals, we therefore limited the maximum injection volume for Juvéderm Voluma® to 200 μL and refrained from shaping. The swelling response and some spontaneous redistribution still took place, but this allowed us to follow the aspect ratio over time despite these unexpected experimental difficulties.

At pre-defined time points the macroscopic dimensions of the implants were assessed with a caliper (day 0, after 3 days, after 3 weeks). For histological evaluation (at 3 weeks for shaped samples, at 6 months and 1 year for unshaped 200 μL samples), mice were sacrificed by intraperitoneal injection of sodium pentobarbital (150mg/kg), followed by transcardial perfusion with 4% paraformaldehyde in PBS. Harvested samples were further immersed for 24 hours in 4% paraformaldehyde at 4°C, washed 3x with PBS and embedded in paraffin following routine procedures. 4 μm slices were stained with hematoxylin/eosin in an automated slide processor (Prisma special stainer, Tissue-Tek; Glas G2 coverslipper from Sakura).

### Magnetic resonance imaging

Magnetic resonance imaging (MRI) images were acquired at 3 days post-implantation on a Varian INOVA console with 14.1 T magnet of 26cm horizontal bore (Magnex Scientific, Abingdon, UK) and a gradient coil (maximum 400 mT/m) with a fix rise time of 120 ms. A home-built single loop surface coil of 24 mm of diameter was used as a transceiver positioned on top of the tissue graft.

During measurements, each adult mouse was anesthetized with 1.5–2% isoflurane in mixture of air and O_2_ (50/50%) and subsequently placed supine within an adapted holder. Body temperature was maintained at 37°C using a thermoregulated water circuit. Monitoring respiration with a respiration cushion was used for triggering during all acquisitions.

Spin echo intensity images were acquired with a repetition time of 1.5s and echo times at 14ms and again at 25ms (the overall intensity of this second echo is used for the images shown). The acquisition matrix was 96×192 pixels for a field of view of 15×25 mm, with 46 slices spaced by 0.05 mm. The acquisition time was on average 12min, with minor differences due to respiratory gating.

### Replication

Unless explicitly stated otherwise, replicated experiments were performed on distinctly prepared biomaterial samples. We further used two 2 separate 1L synthesis batches. Thus the samples contain variability associated with sample preparation, and to some extent chemical synthesis. We used however identical reagent stocks, and so cannot guard against lot-to-lot variation of the chemicals obtained commercially. Supplementary 10 further compares rheological characterization in two distinct geometries.

